# The mating system affects the temperature sensitivity of male and female fertility

**DOI:** 10.1101/2021.06.09.447706

**Authors:** Julian Baur, Dorian Jagusch, Piotr Michalak, Mareike Koppik, David Berger

## Abstract

1. To mitigate effects of climate change it is important to understand species’ responses to increasing temperatures. This has often been done by studying survival or activity at temperature extremes. Before such extremes are reached, however, effects on fertility may already be apparent.
2. Sex differences in the thermal sensitivity of fertility (TSF) could impact species persistence under climate warming because female fertility is typically more limiting to population growth than male fertility. However, little is known about sex differences in TSF.
3. Here we first demonstrate that the mating system can strongly influence TSF using the seed beetle *Callosobruchus maculatus*. We exposed populations carrying artificially induced mutations to two generations of short-term experimental evolution under alternative mating systems, manipulating the opportunity for natural and sexual selection on the mutations. We then measured TSF in males and females subjected to juvenile or adult heat stress.
4. Populations kept under natural and sexual selection had higher fitness, but similar TSF, compared to control populations kept under relaxed selection. However, females had higher TSF than males, and strikingly, this sex difference had increased over only two generations in populations evolving under sexual selection.
5. We hypothesized that an increase in male-induced harm to females during mating had played a central role in driving this evolved sex difference, and indeed, remating under conditions limiting male harassment of females reduced both male and female TSF. Moreover, we show that manipulation of mating system parameters in *C. maculatus* generates intraspecific variation in the sex difference in TSF equal to that found among a diverse set of studies on insects.
6. Our study provides a causal link between the mating system and TSF. Sexual conflict, (re)mating rates, and genetic responses to sexual selection differ among ecological settings, mating systems and species. Our study therefore also provides mechanistic understanding for the variability in previously reported TSFs which can inform future experimental assays and predictions of species responses to climate warming.

## Introduction

To predict evolutionary trajectories of natural populations experiencing warming climates, it is necessary to understand selection on, and the genetic architecture of, traits whose expression depend heavily on temperature (Bubliy et al., 2012; Chevin et al., 2013; Hoffmann & Sgrò, 2011; Walters et al., 2012). Typical estimates of thermal sensitivity describe the temperatures at which individuals fail to maintain basic physiological functions such as controlled locomotion and respiration (Lutterschmidt & Hutchison, 1997). Approaches to predict organismal responses relying exclusively on such measures are however bound to neglect a variety of sublethal effects that will arise at less extreme temperatures, the most important being reductions in fertility (Angilletta, 2009; Chirgwin et al., 2021; Walsh et al., 2019; Parratt et al., 2021; Hoffmann et al., 2013; Kellermann & Heerwaarden, 2019). Indeed, already a slight decrease in fertility can have dramatic effects on population viability (Degioanni et al., 2019) and thermal plasticity in reproductive traits is widespread (Dell et al., 2011; Deutsch et al., 2008; Frazier et al., 2006) and often observable at a significantly lower temperature threshold than responses in viability (Angilletta, 2009; Gerking & Lee, 1983; Hoffmann, 2010; Loisel et al., 2019; van Heerwaarden & Sgrò, 2021). Hence, it is crucial to incorporate estimates of the thermal sensitivity of fertility (from hereon: TSF) into predictions of population persistence (Angilletta, 2009; Parratt et al., 2021; Walsh et al., 2019).

In many sexually reproducing species, population growth is mainly dependent on female fertility (Caswell, 2006). Female TSF may therefore be more consequential for population viability under climate warming, highlighting the need for a more thorough understanding of sex-differences in TSF (Iossa, 2019). For example, if male fertility is more sensitive to elevated temperatures, but assuming some shared genetic basis for TSF in the two sexes, genetic variation with deleterious effects on female TSF could effectively be purged at elevated temperatures while limiting the cost of adaptation (*sensu* Haldane 1957) mainly to males. Such male-biased purging of deleterious alleles affecting TSF could thus pursue with little reduction in population growth, which would aid evolutionary rescue of sexually reproducing species facing warming climates (Godwin et al., 2020; Manning, 1984; Martinossi-Allibert et al., 2019; Plesnar-Bielak et al., 2012; Whitlock & Agrawal, 2009). It has been suggested that male reproduction is more affected by elevated temperature in both endotherms (Hansen, 2009) and ectotherms (David et al., 2005; Jørgensen et al., 2006). However, male and female reproductive physiologies are vastly different (García-Roa et al., 2020; Kodric-Brown & Brown, 1987), questioning whether genetic responses to selection on TSF in one sex would be consequential for TSF in the other. On the other hand, some general buffering mechanisms against elevated temperature, such as antioxidant defences (Dowling & Simmons, 2009) or molecular chaperones aiding protein translation and folding (Feder et al., 2000), are costly to produce and may depend strongly on the overall condition and genetic quality of the individual. Such responses may therefore be much more likely to share a genetic basis between the sexes (Andersson, 1994; Rowe & Houle, 1996; Tomkins et al., 2004).

Sex differences are often rooted in the operation of sexual selection and mating systems, and it is possible that sex-specificity in TSF could trace back to general differences in male and female reproductive physiologies ingrained in the evolution of anisogamy. However, fine-grained variation in sexual selection and mating systems is also likely to play an important role in shaping male and female TSF (García-Roa et al., 2020; Gómez-Llano et al., 2020; Martinossi-Allibert et al., 2019; Pilakouta & Ålund, 2021; Svensson et al., 2020). For example, success under post-copulatory sexual selection (i.e., sperm competition) can depend on both gamete quality (Gage et al., 2004; Hosken et al., 2003; McNamara et al., 2014) and the overall genetic quality of the male (Hosken et al., 2003), suggesting that sexual selection for genetic quality could increase tolerance to thermal stress. However, sperms’ tolerance of oxidative stress, and therefore likely also of high temperature, can be affected by investment in precopulatory traits (Dowling & Simmons, 2009; Helfenstein et al., 2010) and studies have suggested that male gamete quality may trade-off with investment into reproductive competition (Baur & Berger, 2020; Silva et al., 2019). Female reproduction is also sensitive to temperature, especially because egg maturation and oviposition are two highly temperature-dependent processes (Angilletta, 2009; Berger et al., 2008; Kingsolver & Huey, 2008), and it is likely that female TSF could be modulated further by pre- and post-fertilization mating interactions. For example, physical harm inflicted via male harassment of females during copulation, or physiological harm mediated via toxic ejaculate compounds (Arnqvist & Rowe, 2005; Dougherty et al., 2017; Parker, 2006), could increase female TSF if TSF is dependent on the condition of the individual and costly thermal buffering mechanisms. On the other hand, some ejaculate compounds typically have beneficial effects in females (Arnqvist & Nilsson, 2000; Karlsson et al., 1997; Oku et al., 2019; Reinhardt et al., 2009; Savalli & Fox, 1999), and remating could potentially improve male fertility via gamete renewal, suggesting that multiple mating may also have positive effects on TSF. This suggests that sex differences in TSF are bound to vary dynamically with mating system parameters which could have important consequences for evolutionary demography in sexually reproducing species.

Here we explore the role of the mating system in shaping sex differences in TSF in populations of the polyandrous seed beetle *Callosobruchus maculatus*. We first quantified how natural and sexual selection on artificially induced mutations affected short-term evolutionary responses in male and female TSF. This approach was motivated by i) theory often assuming that sexual selection is a more potent force of purifying selection against deleterious genetic variation compared to natural selection (Rowe & Houle 1996, Tomkins et al. 2004, Whitlock & Agrawal 2009) and that purging of deleterious genetic variation is much more efficient via sexual selection in males in *C. maculatus* (Grieshop et al., 2016, 2020), ii) the notion that compensatory physiological responses to temperature stress are costly (Feder et al., 2000), suggesting that TSF may be dependent on the condition and overall genetic quality of the individual, and iii) the observation that elevated temperature can increase the effects of deleterious genetic variation in ectotherms (Berger et al., 2021).

To this end, we induced an appreciable genetic load via mutagenesis in replicate populations that were subsequently propagated for two generations under three alternative experimental evolution regimes: polygamy (imposing natural and sexual selection), enforced monogamy (natural selection only), and relaxed selection (natural and sexual selection removed). Comparisons to the ancestral (non-irradiated) populations allowed us to assess the relative impact of the induced mutations and the extent to which the two mating systems (polygamy and monogamy) had purged the mutations relative to the relaxed selection treatment. Thus, in addition to providing information on sex differences in TSF in *C. maculatus*, this panel of populations allowed us to assess not only sex-specific effects of *de novo* mutations on TSF but also how natural and sexual selection on *de novo* mutations influence sex differences in TSF.

We conducted the experiment using two different types of heat stress. We applied a short-term but high intensity heat shock on adult beetles, reflecting extreme daily maximal temperatures, which are predicted to increase in frequency due to climate change (Johnson et al., 2018). In a parallel experiment using the same populations, we applied long-term heat stress throughout the entire larval development, as may result from increasing variation in average monthly temperatures (Bathiany et al., 2018; Varela et al., 2020). We find no evidence that the genetic load of a population is related to its average TSF. Strikingly, however, experimental evolution under sexual selection increased sex differences in TSF already after two generations. To elucidate the mechanism behind this result we measured TSFs while manipulating male-female (re)mating interactions. Our results show that the mating system can be a key driver of realized TSF in males and females.

## Methods

### Study population

*Callosobruchus maculatus* is a common pest on fabaceous seeds in tropical and subtropical regions. Females cement eggs onto host seeds and the larvae burrow into the seed where they complete their development within 3 weeks under standard laboratory conditions (29° C, 12L:12D light cycle, 55% rel. humidity), on their preferred host, *Vigna unguiculata* (Fox, 1993). Unless otherwise stated, these conditions were also used in the experiments described below. Egg-to-adult survival is above 90% in the populations used here. Adults are facultatively aphageous and start reproducing within hours after emergence. Under laboratory conditions without food or water adult beetles live just over one week with most of the reproduction taking place within the first few days (Fox, 1993). *C. maculatus* has a polyandrous mating system with documented sexual conflict over (re)mating and high remating rates, leading to both pre- and post-copulatory sexual selection on males (Berger et al., 2016; Crudgington & Siva-Jothy, 2000; Eady, 1995; Gay et al., 2009; Hotzy & Arnqvist, 2009). Once a male manages to successfully initiate copulation, spines on its genitalia help to prevent it from being dislodged but at the same time harm the female (Bagchi et al., 2021; Edvardsson & Tregenza, 2005; Rönn et al., 2007; Rönn & Hotzy, 2012). The effects of the genital spines and the harm imposed on the females have been found to correlate with a male’s sperm competitiveness (Hotzy & Arnqvist, 2009) and, in a congener that exhibits similar genital structures, also increase female oviposition rate as a response to genital scarring (Haren et al., 2017). Female reproductive behaviour is further modulated by lifespan-extending nutrients and water (Rönn et al., 2006) and likely also other functional compounds in the male ejaculate (Bayram et al., 2019), suggesting that the male ejaculate can have both positive and negative effects on female fertility (Arnqvist et al., 2004; Yamane et al., 2015).

The stock used for the experimental populations originates from 41 iso-female lines sampled in Lomé, Togo (06°10#N 01°13#E) (see Berger et al., 2014) that were mixed and maintained at large population size (N>300) for roughly 50 generations under standard conditions prior to the start of this experiment.

### Mutagenesis

The stock population was split into three replicate founder populations (Fig. 1a). We first introduced a genetic load in the three founders by exposing male beetles to 25 Gy of γ-radiation over 32 minutes (dose rate: ∼0.79 Gy/min). This dose is known to reduce laboratory fertility (i.e., number of emerging adults) of the parental generation by roughly 70% and that of F1 offspring by roughly 40% (Baur & Berger, 2020; Grieshop et al., 2016). All irradiated males (N = 150 per founder) were virgin and eclosed between 0 and 24 hours prior to irradiation. All males were then mated to a randomly assigned female (enforced monogamy) which was allowed to lay eggs for 48 hours. All egg-laden beans were mixed and distributed into three aliquots marking the starting point for the three different experimental evolution regimes (see below). At this point, all offspring are expected to carry a random set of mutations induced via their fathers. Another 150 control (non-irradiated) males were used to seed the control population for each founder. These control populations were propagated according to standard laboratory protocol.

**Fig. 1:**
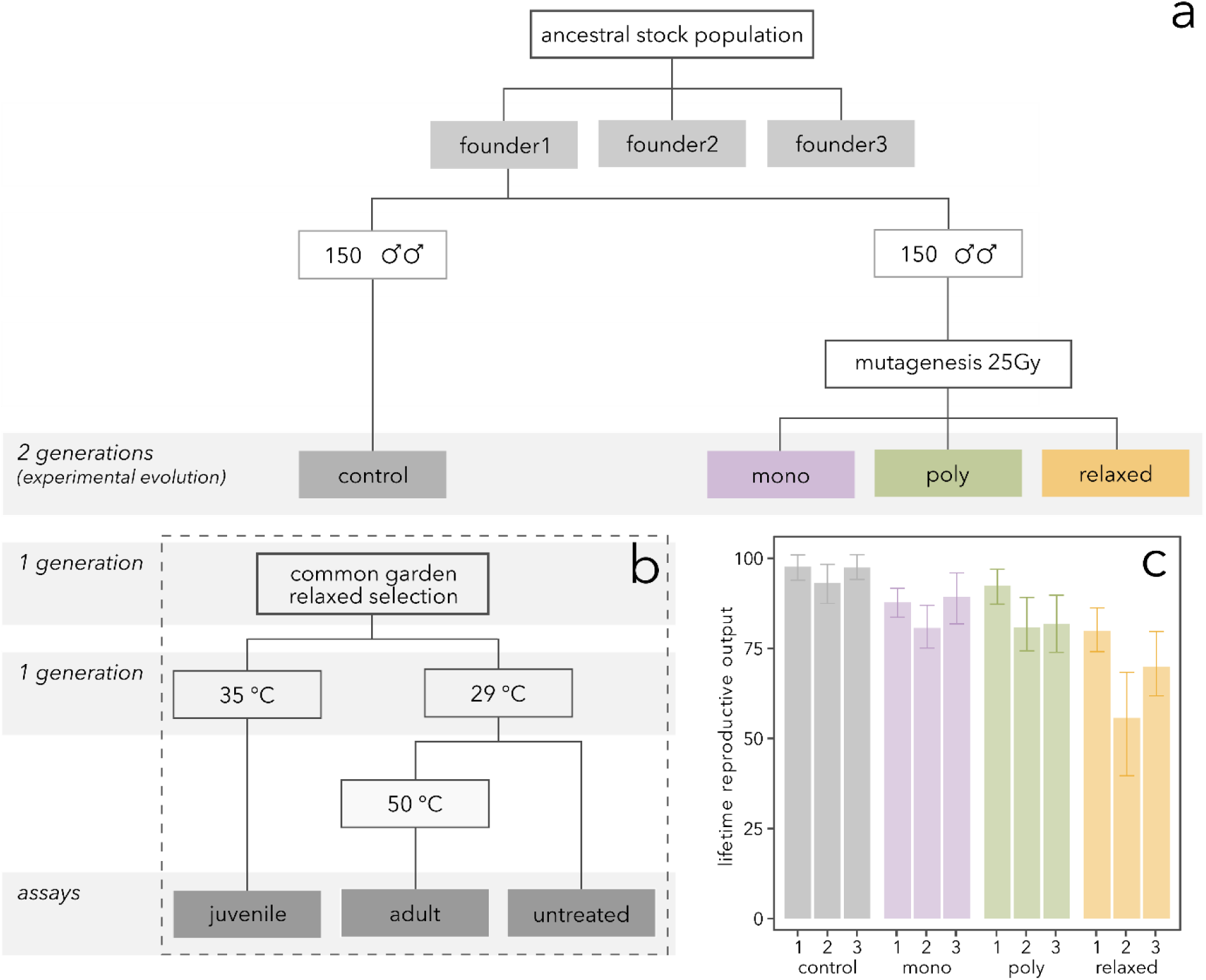
Experimental design used to obtain the two main data sets. a) Founding populations, mutagenesis and experimental evolution. b) Juvenile heat stress (development at 35°C) and adult heat shock (20 min heat shock at 50°C) applied to all 12 populations. c) Lifetime reproductive output of all 12 populations in the benign environment (untreated). Shown are means ± 95% confidence intervals.

### Experimental Evolution regimes

The selection regime protocols (Fig. 1a), outlined below, have previously been described and used in several, more long-term (up to 60 generations), experimental evolution studies in *C. maculatus* and have been shown to result in pronounced sex-specific adaptations (Bagchi et al., 2021; Baur et al., 2019; Baur & Berger, 2020; Martinossi-Allibert et al., 2019).

#### Monogamy

This regime removes sexual selection but applies natural (fecundity) selection on females and males. Within 72 hours after eclosion, 100 virgins of each sex were picked and randomly paired and allowed to mate for 5 hours. During this period, the male and female could freely interact and mate repeatedly. After 5 hours the males were removed, and all females were collected and placed together in a 1-litre jar containing host seeds *ad libitum*. After 48 hours of egg laying the females were removed. To ensure minimal larval competition and viability selection, all populations used in this experiment were provided with beans *ad libitum* for egg deposition (∼4800 black-eyed beans).

#### Polygamy

This regime simulates the natural mating system, including sexual and natural selection on males and females. 100 virgin males and 100 virgin females were picked within 72 hours after eclosion and collected in a 1-litre jar with beans *ad libitum*. The beetles could freely interact, compete, mate and lay eggs for 48 hours, after which all beetles were removed. This mating scheme was also used to propagate the non-irradiated control populations and corresponds to the standard laboratory protocol.

#### Relaxed selection

This regime removes both natural and sexual selection to retain the induced (non-lethal) deleterious mutations in the populations. Within 72 hours after eclosion, 100 virgin females and 100 virgin males were assigned to form random monogamous couples (avoiding inbreeding) as in the monogamous mating regime. Thereafter, males were removed, and each female was provided with beans *ad libitum*. Females laid eggs for 48 hours in isolation, after which all females were removed from the beans. In the next generation, offspring were picked so that each parental couple contributed exactly one female and one male to the next generation.

After two generations of propagation under the respective selection regimes, we applied one generation of common garden relaxed selection to all 12 populations to both counteract potential differences in parental effects brought about by the different evolution regimes, and to prevent further selection against deleterious mutations (Fig. 1a). We then established 30 mating couples per population (total n_family_ = 360). After mating we allowed the female to lay eggs for 48 hours. We then removed the female and evenly split the beans from each female into one half that was subjected to the ***Juvenile heat stress*** treatment (outlined below), and one half that was kept developing at benign 29°C. The beetles developing at 29°C were assigned to undergo the ***Adult heat-shock*** treatment (outlined below) or to remain untreated and serve as control for both heat treatments (Fig. 1a).

### 1. Juvenile heat stress

*1.1.* This experiment was designed to resemble a longer period of elevated temperature as for example a heat wave, which can occur in the months of March and April in Lomé, Togo. Current projections for Lomé predict an increase of the average daily maximal temperature in the months from February to April from 32°C in the late 20^th^ century, up to a maximum of 37.2°C by the end of the 21^st^ century (Varela et al., 2020). Beetles assigned to this treatment developed at an elevated temperature of 35°C throughout their entire larval and pupal stage (ca. 21 days in total). After eclosion, we crossed two treated male and two treated female individuals per family with untreated individuals from other families within the same population, allowing us to estimate the sex-specific fertility loss due to development at 35°C for each of the 12 populations. This resulted in a total of 818 untreated couples, 458 couples with a treated female and 443 couples with a treated male, or roughly 40 couples per treated sex and population. Each couple was provided with beans *ad-libitum* in a 60 mm petri dish and allowed to mate and lay eggs for the rest of their lives. Emerging adult offspring were later counted to obtain the fertility of the couple.

*1.2. Re-mating and male harassment:* We hypothesized that one explanation for our results from the first experiment could be that females developing at stressful temperature might be worse at coping with the harm inflicted by males during mating, but may on the other hand benefit from nutrients in ejaculates. Using the stock population from which the experimental evolution populations were derived, we ran an experiment to tease apart potential effects of remating on female TSF mediated via harmful physical mating interactions and ejaculatory compounds. We exposed 24 hours old virgin heat-treated (developed at 35°C) and untreated (developed at 29°C) females to three male treatments (all males developed at 29° C). Females in the first treatment underwent a single observed mating, after which we removed the male and allowed the female to deposit eggs for the rest of her lifetime. Females undergoing the second treatment were mated once per day to the same male for three consecutive days. After the matings the male was removed. In the third treatment we co-reared the male and female for their entire life, allowing them to interact freely as was the case in the original experiment.

*1.3. Interaction of male and female heat stress*: We also tested whether the cumulative effects of female and male juvenile heat stress act in an additive manner on a couple’s fertility. Beetles of both sexes, originating from the stock population, were reared at benign 29°C and at stressful 35°C, as in the main experiment. Within 24 hours after emergence, we paired a male and female beetle of which either the male, the female, both sexes, or none of the sexes, had developed at elevated temperature. Males and females were co-reared for their entire life, and we counted the couple’s reproductive output.

### 2. Adult heat shock

*2.1.* This experiment was designed to simulate a short-term heat extreme as occurring in the form of extreme daily maxima. We chose an exposure intensity of 50°C for 20 minutes because a pilot experiment showed that ca. 50% of the beetles are knocked-out in the process (no more perceptible movement), but at the same time it remains in the range that we consider ecologically relevant as such temperatures are likely to be reached in sun exposed microclimates within the range of this species (Deutscher Wetterdienst, n.d.). From the beetles that developed at benign temperature, we randomly picked three adult female and three adult male beetles per family, resulting in 818 untreated couples, 500 couples with a treated female and 540 couples with a treated male. We put beetles in a perforated 0.5 ml Eppendorf tube placed in a closed 200 mm petri dish on a heating plate set to 50°C. The air temperature inside the upper part of the petri dish was also monitored and remained constant at 43°C for the duration of the treatment. We paired all heat-exposed beetles with an untreated individual of opposite sex from a different family, but of the same population. Each couple was provided with *ad-libitum* beans in a 60 mm petri dish and lifetime reproductive output was recorded.

*2.2. Male recovery:* Using the stock population, we investigated if recovery of the male after heat shock may have shaped the observed sex differences in TSF in the main experiment outlined above. We allowed one group of males to mate 2 hours after the heat shock treatment. Beetles of this group were then allowed to mate again 26 hours after heat shock. The second group of male beetles were only allowed to mate once, after 26 hours. Beetles assigned to the untreated control group were subdivided into the same two groups to control for a possible decline in fertility due to repeated mating, although this has been shown to be minimal in *C. maculatus* (Rönn et al., 2008). This allowed us to independently estimate effects of recovery over time and recovery through mating, causing ejaculate replacement, on male TSF.

### Statistical analysis

All analyses were executed in R (R Core Team, 2020). We fitted generalized linear mixed models assuming a Poisson distributed response using the package lme4 (Bates et al., 2015, p. 4), unless stated otherwise. The R package ggplot2 (Wickham, 2016) was used for graphical illustration. P-values were calculated using the package car (Fox & Weisberg, 2019) using type-II sums of squares. Planned post-hoc comparisons, applying Tukey correction, were conducted using the package emmeans (Lenth, 2020).

*1.1 & 2.1 Sex-specific TSF in evolution regimes*. Offspring number was used as the response while evolution regime and treatment (male stress, female stress or untreated), as well as their interaction, were added as fixed effects. We also added experimental block and the identity of the experimenter counting offspring as additional terms. Population replicate crossed with treatment, as well as dam and sire effects, were included as random effects. Additionally, an observation level random effect (OLRE) was included to control for over-dispersion.

*1.2. Re-mating and male harassment*: We analysed main effects of female development treatment (29°C/35°C), mating (single/multiple) and cohabitation (isolated/cohabiting), as well as two-way interactions between development treatment and mating and cohabitation, respectively. Experimental date was added as an additional main effect. We assumed Poisson-distributed errors while correcting for overdispersion via the quasi-extension in both models.

*1.3. Interaction of male and female heat stress*: We analysed effects of female development treatment (29°C/35°C), male development treatment (29°C/35°C) and their interaction as fixed factors assuming Poisson-distributed errors.

*2.2 Male recovery*: We analysed effects of a recovery treatment (mated 2 or 26 hours after the heat shock) with and without remating separately. To test for the effect of remating after heat shock, we ran a generalized linear mixed model with a Poisson response including treatment (untreated, heat shocked), mating number (one (2h) or two (26h)) and their interaction as fixed effects, and male identity as well as an OLRE as a random effect. To analyses the effect of recovery without mating, we used the same model type and structure but without the male identity and the interaction term because males only mated once. In this model treatment included untreated, heat shock and 2 hours recovery, and heat shock and 26 hours recovery.

## Results

There were pronounced fertility differences between irradiated and control populations when assayed only in the benign environment (i.e., untreated beetles), illustrating that mutagenesis had induced a sizeable genetic load (Χ^2^= 60.88, df = 3, p < 0.001) (Fig. 1c, Supplementary table 1). Pairwise post-hoc comparisons revealed that the regime under relaxed selection carried a larger genetic load than the monogamy and the polygamy regime, demonstrating efficient purging of deleterious mutation during experimental evolution (Tukey_relax-poly_: z = -4.52, p < 0.001; Tukey_relax-mono_: z = -5.16, p = p < 0.001). There was no difference in fertility between the monogamy and polygamy regime (Tukey_poly-mono_: z = -0.68, p = 0.91). All pairwise contrasts in supplementary table S1.

### Juvenile heat stress

*1.1.* Elevated temperature during juvenile development strongly reduced the reproductive output of adults (Χ^2^ = 179.07, df = 2, p < 0.001). In the control populations, a female developing at 35°C showed an average reduction in fertility of 32% (31 fewer offspring) compared to an untreated female, while male fertility was reduced by 22% (21 fewer offspring). To investigate if the induced mutations affected TSF, we first compared the effect of heat stress in the control regime and the regime evolving under relaxed selection (containing the largest genetic load; Fig. 1c). We found no evidence that genetic load affected TSF (Χ^2^ = 3.75, df = 2, p = 0.15). Strikingly, however, sex differences in TSF depended on selection regime (sex:regime; Χ^2^= 14.04, df = 6, p = 0.029) (Fig. 2a, b). There were clear reductions in fertility via heat stress in both sexes in all but the polygamy regime, where exposed males showed no statistically significant fertility loss (Fig. 2a, Supplementary table S2). To directly assess the effect of sexual selection on TSF we compared the monogamy (natural selection) and polygamy (natural + sexual selection) regimes. This analysis confirmed the results of the global model (sex:regime; Χ^2^ = 6.96, df = 2, p = 0.030, Fig 2b). Heat-treated females from the polygamy regime produced significantly fewer offspring than heat-treated females from the monogamy regime (Tukey_poly_female35-mono_female35_: z = -2.08, p = 0.04), while heat-treated polygamy males instead tended to produce more offspring than heat-treated monogamy males, although this effect was not statistically significant (Tukey_poly_male35-mono_male35_: z = 1.41, p = 0.16). Full model specification and output in supplementary material S3.

**Fig. 2:**
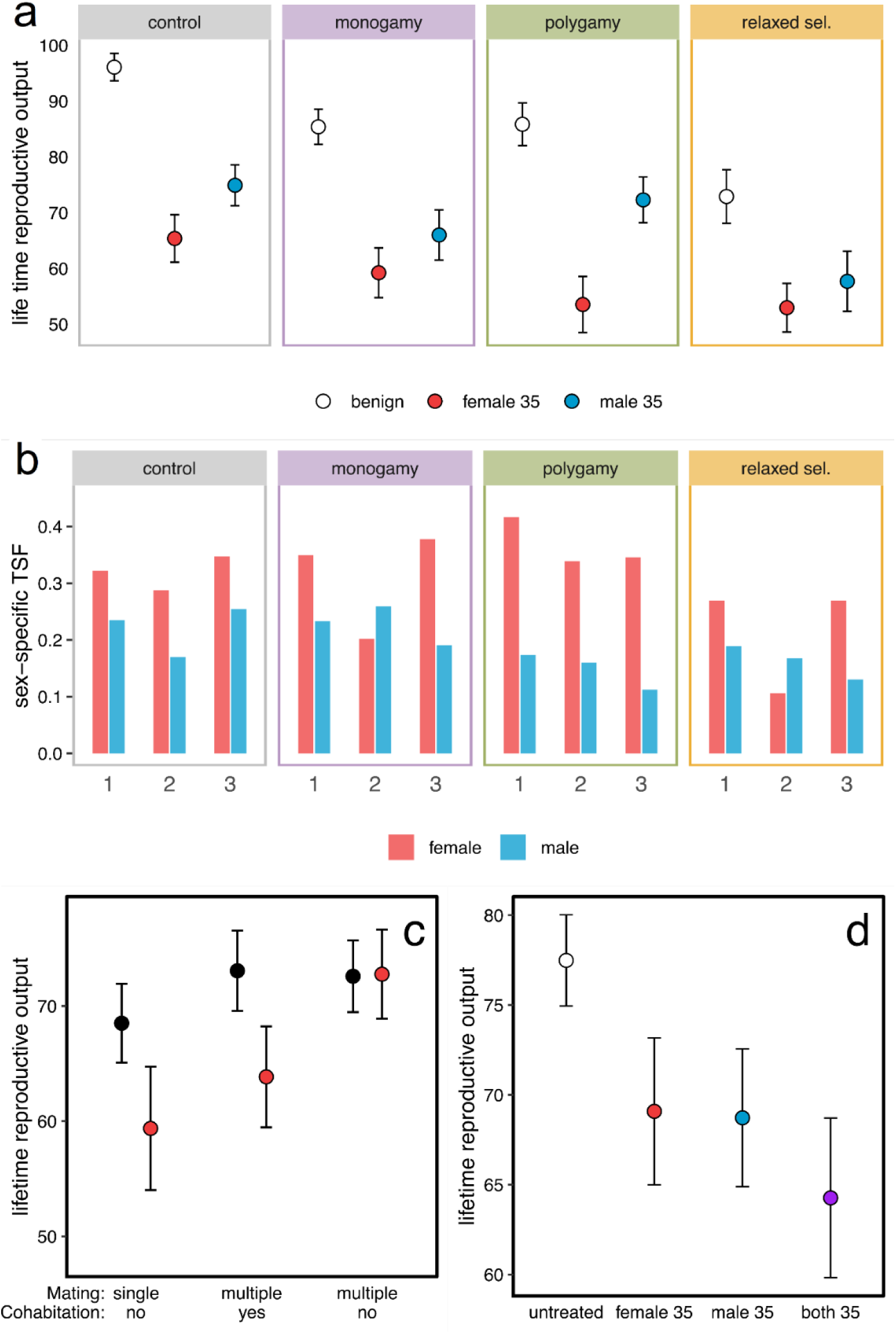
Sex differences in TSF under juvenile heat stress (experiments *1.1-1.3*) a) Lifetime reproductive output of couples with either the female (red symbols), the male (blue symbols), or no parent (open symbols) developing at elevated temperature. b) Relative loss in fertility (1-stressed/control) per population. Sex differences in TSF in all three blocks are greater for polygamy populations compared to all other regimes. c) Lifetime reproductive output of female beetles developing at benign (black symbols) or stressful (red symbols) temperature. d) Lifetime reproductive output of pairs in which no parent (white), the female (red), the male (blue), or both parents (purple) were exposed to juvenile heat stress. Panels a, c and d show means ± 95% confidence intervals.

*1.2.* To gain more insights into the underlying mechanisms responsible for the evolved sex difference in TSF, we ran an additional experiment on the stock population to investigate the role of repeated mating and male harassment on female TSF. Females exposed to juvenile heat stress suffered more under cohabitation with a male than females developing at benign temperature (temperature:cohabitation; Χ^2^ = 4.53, df = 1, p = 0.033, Fig 2c), which could in part explain the evolved increase in female-bias of TSF observed in the polygamy regime, if polygamy males were more persistent during mating. Interestingly, there was also a positive effect of re-mating, and this effect was more beneficial in females developing at elevated temperature (temperature:mating; Χ^2^ = 4.99, df = 1, p = 0.025). Crucially, however, the beneficial effect of re-mating depended strongly on the exclusion of the male between matings (Tukey_single_35 vs. cohabitation_35_: z = -1.62, p = 0.24, Tukey_single_35 vs. remated_35_: z = -4.345, p < 0.0001, Fig. 2c); and strikingly, re-mating with experimental exclusion of the male between matings sufficed to completely thwart the negative effect of developmental temperature stress in females (Tukey_remated_29 vs. remated_35_: z = -0.40, p = 0.69). Hence, changes in the relative costs and benefits of multiple mating between the polygamy and monogamy regime is likely a driver of the evolved sex difference in TSF.

*1.3.* We also explored whether the effect of males on female TSF was dependent on whether males had also been exposed to heat. As expected, we found strong effects of both male and female developmental heat stress (Female: Χ^2^= 11.98, df = 1, p < 0.001; Male: Χ^2^ = 10.75, df = 1, p = 0.001) (Fig. 2d). The interaction between female and male heat stress, however, was non-significant, suggesting that the effects of female and male juvenile heat stress on TSF are mostly additive. This result might be explained by heat stress reducing the underlying male components with antagonistic effects on female TSF (level of male harm and beneficial ejaculate compounds) to similar extent.

### Effects of adult heat shock

*2.1.* The adult heat shock treatment led to an overall loss of fertility (Χ^2^= 17.17, df = 2, p < 0.001) even though its impact was much weaker compared to the impact of juvenile heat stress (average fertility loss for females was 10.2% and for males 4.2%). The effect of heat shock was significantly stronger in females (Tukey_female – male_: z = -2.38, p = 0.045) and, in fact, not statistically detectable in males (Tukey_untreated – male_: z = 2.08, p = 0.09) (Fig. 3a, Supplementary table S2). The effect of heat shock was generally too weak to be detected when analysing subsets of the data (results not shown), resulting in neither the induced genetic load (regime_control vs. relaxed_:treated sex: Χ^2^ = 0.44, df = 2, p = 0.88) nor sexual selection (regime_polygamy vs. monogamy_:treated sex: Χ^2^ = 0.43, df = 2, p = 0.81) having a statistically significant effect on TSF. Full model specification and output in supplementary material S4.

**Fig. 3:**
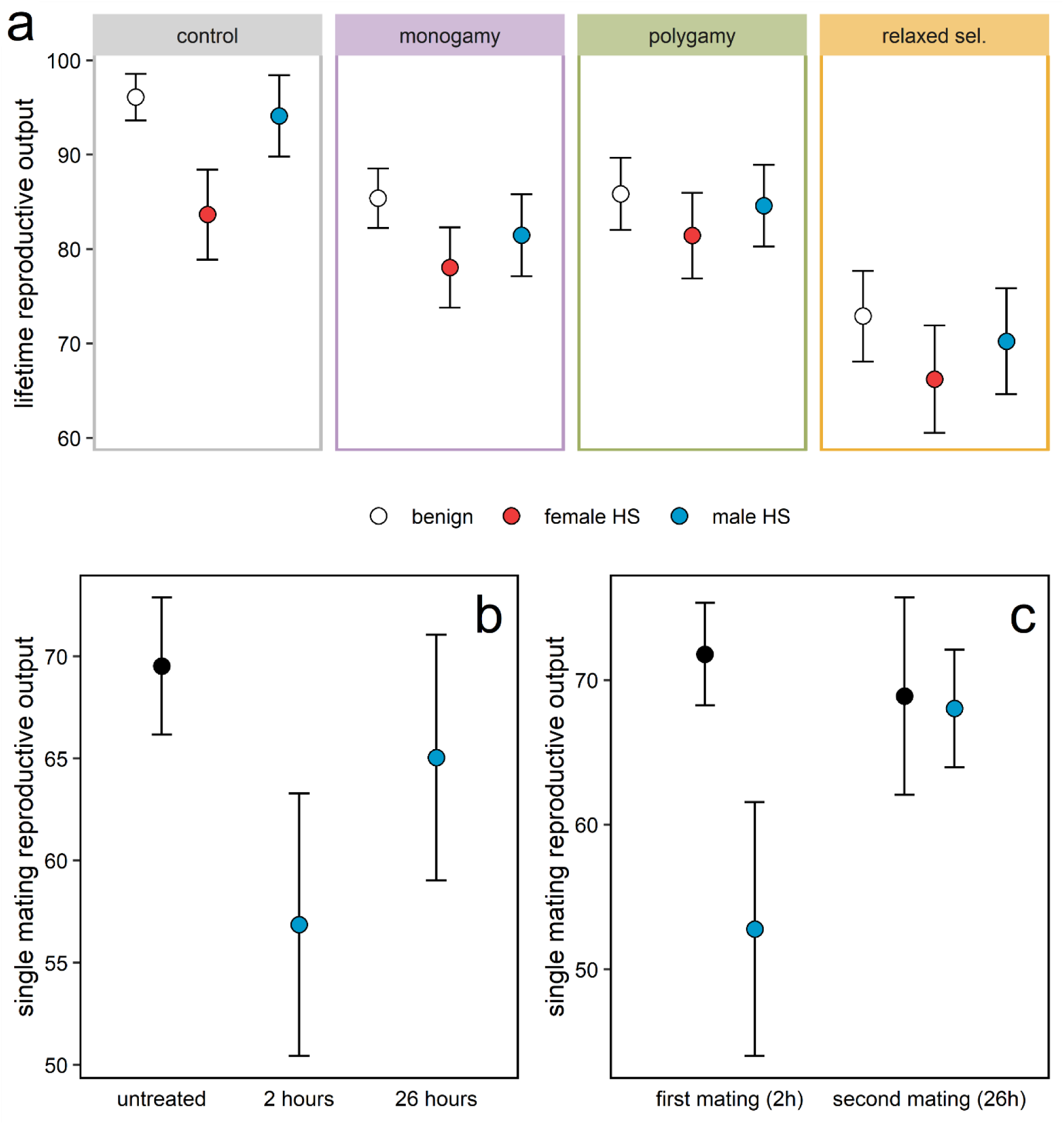
Sex differences in TSF under adult heat shock (Experiments *2.1 & 2.2*) a) Lifetime reproductive output depending on the sex that underwent adult heat shock (HS). b) Recovery of male fertility after adult heat shock. c) Lifetime reproductive output of untreated (black) and heat-shocked (blue) males that were mated to a virgin female 2 hours after the treatment and again, to a second female, 26 hours after the treatment. All panels show means ± 95% confidence intervals.

*2.2.* To elucidate underlying mechanisms explaining sex differences in TSF under adult heat shock, we analysed effects of male recovery, in terms of both time and remating (inducing ejaculate renewal) after heat shock. Our data show that male beetles can recover almost completely from the applied heat shock treatment within a 26-hour recovery period. Males showed strong TSF, signified by an 18% reduction in fertility compared to the untreated control group, when mating within two hours after heat shock (Tukey_untreated – 2 hours_: z = 3.55, p = 0.001, Fig. 3b). If males were given 26 hours of recovery in isolation, however, no significant effect on fertility could be found (Tukey_untreated – 26 hours_: z = 1.14, p = 0.49) (Fig. 3b). Similarly, there was no reduction in fertility in beetles that mated a second time 26 hours after the treatment (Tukey_untreated 2 hours – 2 hours_: z = 0.64, p = 0.003; Tukey_untreated 26 hours – 26 hours_: z = 0.006, p = 1) (Fig. 3c). Recovery with or without remating showed similar effects on TSF (Tukey_heat 26 hours mate – heat 26 hours isolated_: z = 0.33, p = 0.74), suggesting that timing is crucial when assessing TSF, and that realized sex differences in TSFs in natural populations are state-dependent properties of mating system and ecology.

### Comparing intra- and interspecific variation in sex differences in TSF

To put our results into perspective, we performed a (non-exhaustive) literature search for studies on other insects that had estimated effects of heat stress on both male and female fertility (summarized in Supplement S5 and S6). This allowed us to calculate and compare standardized estimates of sex differences in TSF (Fig. 4). The variability in this estimate obtained by manipulating mating system parameters and the timing of heat stress relative to (re)mating in our study roughly corresponds to that reported between species in previously published studies, demonstrating that the mating system can be a main determinate of sex differences in TSF. Moreover, in contrast to occasional claims of male biased TSF, there is no such consistent bias in the reviewed studies on insects estimating male and female TSF under the same experimental conditions.

**Fig. 4:**
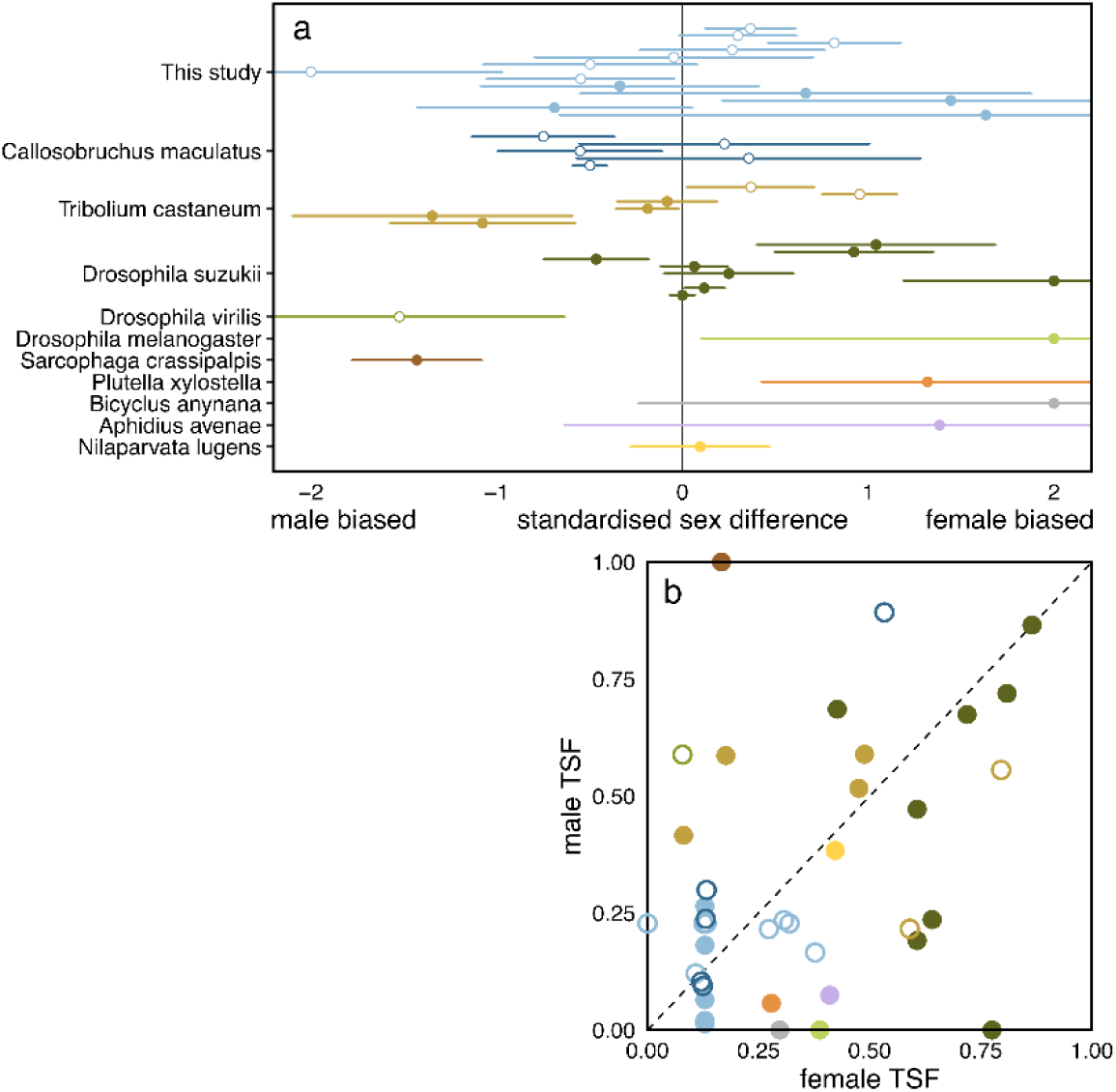
Intraspecific variation in the sex difference in TSF generated through manipulation of the mating system, timing of heat stress relative to (re)mating in this study, compared to estimates from other studies on insects. Open symbols represent heat treatments applied at a juvenile stage while closed symbols represent heat treatments applied during the adult stage. a) A standardized measure of the sex difference in TSF was calculated as: SD_TSF_ = 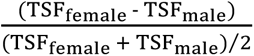, where TSF is the relative fertility loss due to heat stress (1-benign/stress). A given SD_TSF_ from this study was derived by first calculating the TSF of one sex in a given experimental condition (e.g., TSF_female_ with remating but no cohabitation from experiment *1.2*) and then always using the TSF observed in one of the two main experiments (*1.1* for juvenile stress and *2.1* for adult stress) for the opposite sex from control populations as comparison (e.g., TSF_male_ from experiment *1.1*). 95% confidence intervals were calculated through propagation of the uncertainty reported for measures of reproductive output within the respective studies. b) Comparison of male and female TSF for the studies presented in panel a. Further details, including inclusion criteria for reviewed studies and a table of all values, are presented in Supplementary material S5 & S6.

## Discussion

In this study we have demonstrated that the mating system can affect sex differences in TSF using the seed beetle *C. maculatus*. Strikingly, sexual selection on induced mutations over only two generations of experimental evolution led to increased female-bias in TSF in polygamous populations experiencing developmental heat stress. Male harassment aggravated the negative effects of heat stress on females, suggesting that increased male harassment might explain the increased female-bias in TSF in polygamous experimental populations. In *C. maculatus,* sexual selection in males is more than three times as effective at purging deleterious alleles compared to fecundity selection on females under semi-natural laboratory setting, as used here (Grieshop et al. 2016, Grieshop et al. 2021). One plausible mechanism behind the result is therefore that sexual selection in the polygamous mating regime led to more efficient purging of alleles with deleterious effects on male mating success, relative to purging of alleles with deleterious effects on female viability and fertility, potentially shifting the balance between male persistence and female resistance during (re)mating interactions. Male-biased selection on deleterious alleles can also improve population fitness by sparing females the cost of adaptation (Manning 1984, Agrawal 2001, Siller 2001, Agrawal & Whitock 2009), if some of the deleterious alleles in males also have deleterious effects in females (Andersson 1994, Rowe & Houle 1996, Chippindale et al. 2001, Tomkins 2004, Bonduriansky & Chenoweth 2009), for which there is evidence in *C. maculatus* (Grieshop et al. 2021). However, once males and females in the polygamous populations engaged in mating interactions, the heightened genetic quality of male genotypes evolving under strong sexual selection may have resulted in increased male harassment of females, and the negative effects of this sexual conflict may have been exposed under female heat stress. In nature, the relative extent of this negative effect should strongly depend on ecological settings and population densities modulating the degrees of conflict (Arbuthnott et al., 2014; Gomez-Llano et al., 2018; MacPherson et al., 2018; Yun et al., 2017). We also note that our comparison of monogamous and polygamous populations evolving from inflated levels of mutational variation does not describe a natural scenario of long-term evolution in populations under mutation-selection balance. Instead, our approach was designed to reveal how (sex-specific) natural and sexual selection can act on genetic variation to shape TSF. Hence, our study provides a proof-of-principle for a direct link between the mating system and sex differences in TSF.

At present, the scant literature available seems to suggest that male reproduction is more sensitive to heat stress than female reproduction (David et al., 2005; Porcelli et al., 2017; Sales et al., 2018), and male fertility has also been demonstrated to be very temperature-sensitive in *C. maculatus*. Our data show that *C. maculatus* females can in fact be more strongly affected by heat stress, and that the realized TSF in males and females can be highly contingent on the experimental design (see also Terblanche et al., 2007) and mating system parameters such as the extent of sexual conflict and remating rates. For example, the adult heat shock treatment resulted in relatively weak effects on fertility, but with significant female-bias in TSF. However, our additional experiment showed that males fully recovered from heat shock within only 26 hours, implying that the sex-bias in TSF in adults is likely to change throughout life following heat exposure. In the case of juvenile heat stress, male harassment aggravated effects of heat stress on female fertility, while repeated mating instead had positive effects on both male and female TSF. However, the size of the nuptial gift provided by male *C. maculatus* has been found to decrease with temperature, suggesting that this male compensatory effect may diminish when also males are heat stressed (Fox et al., 2006). Other studies on fruit flies (García-Roa et al., 2019, 2020) and *C. maculatus* (Martinossi-Allibert et al., 2019), conclude that heat stress generally reduces the female fertility cost of male cohabitation. In both these studies, the impact of sexual conflict was assessed directly in the stressful environment when both sexes were stressed, while in the present study the effects of mating interactions on fertility were measured in a benign environment after heat stress had been applied to one, or both sexes. Collectively, this limited set of studies suggest that there is a multitude of ways that temperature can modulate the consequences of sexual selection and conflict (Garcia-Roa et al. 2020), and conversely, that sexual selection and conflict can shape sensitivity to temperature (Martinossi-Allibert et al. 2019). Depending on population density, mating system, and heat stress characteristics, laboratory experiments may thus lead to erroneous estimates of TSFs, and in extension, misjudgements of the threat on population growth imposed by climate warming, even when efforts are made to measure TSF sex-specifically. Indeed, our comparison of variation in sex differences in TSF generated by mating system parameters in our study, to that reported between species (Fig. 4), suggests that TSF is a highly dynamic property that responds to population structure and ecological changes.

Directly comparing the effects of our two heat stress treatments is difficult as they were applied with different intensities over different time frames and life stages. Nevertheless, juvenile heat stress is known to affect the development of reproductive organs and result in reduced sperm numbers in insects (Chirault et al., 2015; Kirk Green et al., 2019; Nguyen et al., 2013; Vasudeva et al., 2014). Vasudeva, Deeming and Eady (2014), found not only a decrease in sperm number but also a reduction in relative testis size by almost 25% in *C. maculatus* males exposed to similar juvenile heat stress as in our experiment. In a later study, the same authors determined the first 20% of larval development to be the most temperature sensitive period of testis development (Vasudeva et al., 2021). A recent study using the flour beetle *Tribolium castaneum* found that the most sensitive phase of testis development is likely during the pupal stage and that testis size can be almost complete recovered in males exposed to heat stress at an immature adult stage (Sales et al., 2021). Together, these studies suggest that there are several time points with heightened temperature sensitivity throughout male reproductive development. Heat shock treatments applied on adults have also been found to decrease numbers of transferred sperm and reduce fertility (Chevrier et al., 2019; Sales et al., 2018, 2021), but considering the data presented here and in Sales et al. (2021), such effects may be reversible in most cases. Similar changes in the morphology of female reproductive organs (i.e., smaller ovaries) combined with a strong reduction in egg number have also been reported for flies of the species *Drosophila suzukii* developing at elevated temperature (Kirk Green et al., 2019). However, little is known about the ability of female reproduction to recover from heat stress. Data from an experiment exposing newly emerged cotton bollworm females, *Helicoverpa armigera*, to a range of heat shock treatments shows a postponement of peak reproduction correlated to the treatment intensity, suggesting some recovery processes taking place between the heat shock event and the onset of reproduction (Mironidis & Savopoulou-Soultani, 2010). In summary, this suggest that, at least in holometabolous insects, heat stress during development can cause an impairment of reproductive organs which is only reversible given a considerable amount of recovery time (if at all), while heat shock experienced at the adult life stage might be reversible on shorter time scales. Importantly, however, we also show that strategies such as remating or postponement of reproduction may mitigate the impact of heat stress experienced both early and late in life.

Inducible compensatory responses that buffer the effects of heat stress are costly and may therefore depend on the genetic quality of the organism. Moreover, it has recently been shown that elevated temperatures can aggravate the deleterious effect of mutations (Berger et al., 2021). We therefore predicted that populations with larger genetic loads might show increased TSF but found no support for this in our data. The model applied by Berger et al. (2021) predicts that temperature-dependent increases in mutational effects stem from reversable misfolding of proteins at high temperature. It is possible that such effects were no longer apparent following heat stress in our experiment as individuals were shifted back to benign temperature (i.e., temperature-sensitive mutants either died during development, or survived and got “rescued” by being placed at benign temperature). Indeed, individuals surviving short term heat stress may even elicit compensatory stress responses that mitigate deleterious effects of mutations (Casanueva et al., 2012). Additionally, as we here studied temperature effects in the adult stage, where realized TSFs are consequences of mating interactions, it is possible that a weakening of sexually antagonistic interactions in populations with large genetic loads (and low-condition individuals) may have contributed to mitigating the detrimental effects of temperature. Indeed, a general tenet highlighted throughout this study how frequency-dependent processes in general (Bolnick et al., 2011; Brady et al., 2019; Dall et al., 2012; Svensson & Connallon, 2019), and sexual selection in particular (Chenoweth et al., 2015; García-Roa et al., 2020; MacPherson et al., 2018; Martinossi-Allibert et al., 2019; Martinossi-Allibert et al., 2019; Rankin et al., 2011; Yun et al., 2017) may affect population vulnerability and adaptation to abiotic factors.

## Acknowledgements

We would like to thank Elena Mondino for help with the illustration of the experimental design, Johanna Liljestrand Rönn for help in the laboratory, and Bo Stenerlöw and Olga Vorontsova for help with the irradiation. This research was funded by grants from the Swedish research council (VR: 2015-05223 and 2019-05024) and Carl Tryggers Stiftelse (CTS:18:32) to DB.

## Author contributions

J.B., D.B. and M.K. conceived the ideas and designed the experiments; J.B., D.J., P.M.M. and M.K. collected the data; J.B. and D.B. analysed the data; J.B. and D.B. led the writing of the manuscript. J.B., D.B. and M.K. contributed critically to the drafts and all authors gave final approval for publication.

## Data availability

All data presented in this manuscript will be archived and made publicly available on Dryad upon acceptance.

## Supplementary material

**Supplement S1:** Table of all pairwise contrasts between regimes for offspring numbers at benign temperature.

| Contrast | Estimate | Std.Error | z | P-value |
| --- | --- | --- | --- | --- |
| monogamy - control | -0.122 | 0.045 | -2.713 | **0.034** |
| polygamy - control | -0.154 | 0.046 | -3.369 | **0.004** |
| relaxed - control | -0.392 | 0.050 | -7.778 | **<0.001** |
| polygamy - monogamy | -0.032 | 0.048 | -0.675 | 0.906 |
| relaxed - monogamy | -0.270 | 0.052 | -5.159 | **<0.001** |
| relaxed - polygamy | -0.238 | 0.053 | -4.523 | **<0.001** |

**Supplement S2:**
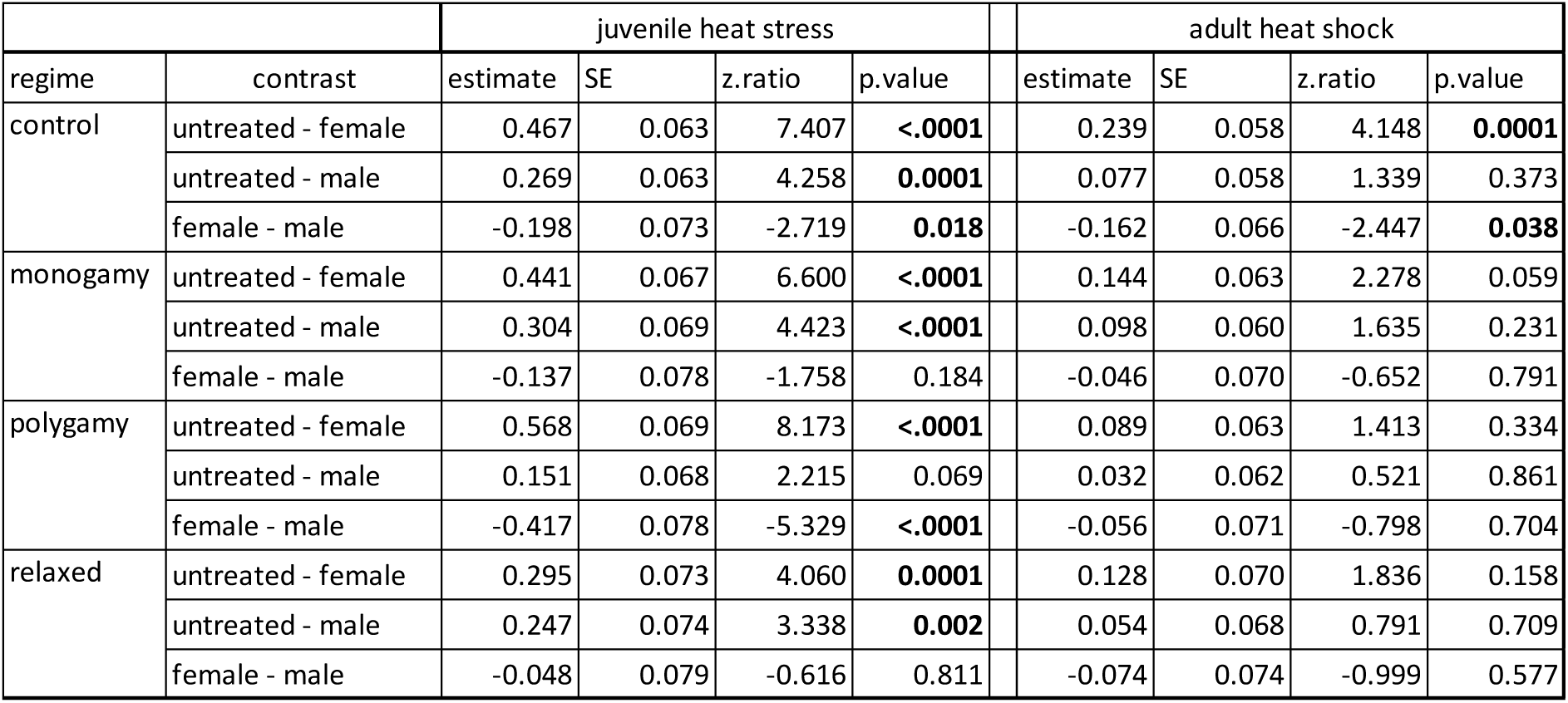
Pairwise comparisons between treatments (untreated, stressed female, stressed male) within selection regimes for both juvenile heat stress and adult heat shock main datasets (experiments 1.1 and 2.2). Marginal means were obtained by averaging over blocks and experimenter. All p-values are Tukey corrected.

**Supplement S3:** Details global model – juvenile heat stress glmer(offspring ∼ regime*treatment + block + counter + (1|line) + (1|treatment:line) + (1|familyID_female) + (1|familyID_male) + (1|OLRE_ID), family=“poisson”, data = data, control=glmerControl(optimizer=“bobyqa”,optCtrl=list(maxfun=100000))) Sample size: 1719, groups: OLRE_ID, 1719; familyID_male, 344; familyID_female, 341; treatment:line, 36; line, 12

Random effects:
| Groups | Variance | Std.Dev. |
| --- | --- | --- |
| OLRE_ID | 2.94E-01 | 5.42E-01 |
| familyID_male | 1.76E-02 | 1.33E-01 |
| familyID_female | 8.32E-03 | 9.12E-02 |
| treatment:line | 1.32E-15 | 3.63E-08 |
| line | 0.00E+00 | 0.00E+00 |

Fixed effects:
|  | Estimate | Std.Err. | z | Pr(>\|z\|) |
| --- | --- | --- | --- | --- |
| Intercept (ctrl, benign temp.) | 4.5983 | 0.04607 | 99.81 | **<0.001** |
| regimemono | -0.12952 | 0.05794 | -2.235 | **0.02539** |
| regimepoly | -0.16538 | 0.05886 | -2.81 | **0.00496** |
| regimerelax | -0.41747 | 0.06406 | -6.517 | **<0.001** |
| treatedFemale | -0.46662 | 0.06299 | -7.407 | **<0.001** |
| treatedMale | -0.2685 | 0.06305 | -4.258 | **<0.001** |
| block2 | -0.18958 | 0.03962 | -4.785 | **<0.001** |
| block3 | -0.08568 | 0.04038 | -2.122 | **0.03383** |
| counter2 | 0.01375 | 0.02828 | 0.486 | 0.62678 |
| regimemono:treatedFemale | 0.02598 | 0.09176 | 0.283 | 0.77705 |
| regimepoly:treatedFemale | -0.10092 | 0.09367 | -1.077 | 0.28127 |
| regimerelax:treatedFemale | 0.17153 | 0.09606 | 1.786 | 0.07416 |
| regimemono:treatedMale | -0.03515 | 0.09322 | -0.377 | 0.70617 |
| regimepoly:treatedMale | 0.11752 | 0.09282 | 1.266 | 0.20548 |
| regimerelax:treatedMale | 0.02184 | 0.097 | 0.225 | 0.82189 |

Anova table (type II SS):
|  | X2 | Df | Pr(>X2) |
| --- | --- | --- | --- |
| regime | 60.5142 | 3 | **<0.001** |
| treatment | 179.0658 | 2 | **<0.001** |
| block | 22.9029 | 2 | **<0.001** |
| counter | 0.2365 | 1 | 0.62678 |
| regime:treatment | 14.0398 | 6 | **0.02919** |

Random effects:

**Supplement S4:** Details global model – adult heat shock glmer(offspring ∼ regime*treatment + block*treatment + counter + (1|line) + (1|treatment:line) + (1|familyID_female) + (1|familyID_male) + (1|OLRE_ID), family=“poisson”, data = data, control=glmerControl(optimizer=“bobyqa”,optCtrl=list(maxfun=100000))) Sample size: 1858, groups: OLRE_ID, 1858; familyID_male, 343; familyID_female, 343; treatment:line, 36; line, 12

| Groups | Variance | Std.Dev. |
| --- | --- | --- |
| OLRE_ID | 2.67E-01 | 5.17E-01 |
| familyID_male | 2.29E-02 | 1.51E-01 |
| familyID_female | 8.62E-03 | 9.28E-02 |
| treatment:line | 0.00E+00 | 0.00E+00 |
| line | 0.00E+00 | 0.00E+00 |

Fixed effects:
|  | Estimate | Std.Err. | z | Pr(>\|z\|) |
| --- | --- | --- | --- | --- |
| Intercept (ctrl, benign temp.) | 4.59271 | 0.04831 | 95.067 | **<0.001** |
| regimemono | -0.12807 | 0.05713 | -2.242 | **0.02498** |
| regimepoly | -0.16222 | 0.05798 | -2.798 | **0.00515** |
| regimerelax | -0.40836 | 0.06316 | -6.465 | **<0.001** |
| treatedFemale | -0.14945 | 0.0695 | -2.15 | **0.03152** |
| treatedMale | -0.05093 | 0.06956 | -0.732 | 0.46409 |
| block2 | -0.20795 | 0.05093 | -4.083 | **<0.001** |
| block3 | -0.07442 | 0.05336 | -1.395 | 0.16309 |
| counter2 | 0.03837 | 0.02666 | 1.439 | 0.15014 |
| regimemono:treatedFemale | 0.09522 | 0.08499 | 1.12 | 0.26255 |
| regimepoly:treatedFemale | 0.15054 | 0.08471 | 1.777 | 0.07554 |
| regimerelax:treatedFemale | 0.11093 | 0.09022 | 1.229 | 0.21889 |
| regimemono:treatedMale | -0.02127 | 0.08308 | -0.256 | 0.79788 |
| regimepoly:treatedMale | 0.04498 | 0.0842 | 0.534 | 0.59318 |
| regimerelax:treatedMale | 0.02312 | 0.08897 | 0.26 | 0.795 |
| treatedFemale:block2 | 0.05539 | 0.07281 | 0.761 | 0.44682 |
| treatedMale:block2 | 0.07261 | 0.07233 | 1.004 | 0.31539 |
| treatedFemale:block3 | -0.32439 | 0.07928 | -4.091 | **<0.001** |
| treatedMale:block3 | -0.15126 | 0.07624 | -1.984 | **0.04724** |

Anova table (type II SS):
| | $\chi^2$ | Df | Pr(>X2) |
| --- | --- | --- | --- |
| regime | 68.2803 | 3 | **<0.001** |
| treatment | 17.1739 | 2 | **<0.001** |
| block | 31.7977 | 2 | **<0.001** |
| counter | 2.0708 | 1 | 0.1501429 |
| regime:treatment | 4.0144 | 6 | 0.6747337 |
| treatment:block | 25.3058 | 4 | **<0.001** |

**Supplement S5:** Literature review

To put the variation in the sex-specificity of TSF obtained across the various data sets presented in our study into perspective, we performed a non-exhaustive literature search for studies reporting sex-specific effects of heat stress on fertility measured for both sexes within the same experiment applying the same heat stress treatment. The literature search was executed between the 3.5.2021 and the 22.5.2021 using Google Scholar. We considered the results of a search for titles using the key words “fertility”, “heat”, “reproduction”, “sex”, “stress”, and “temperature”, as well as literature cited by papers matching the initial search. Note that the literature search was non-exhaustive, but that we included all studies we could find that had measured fertility for both sexes in terms of reproductive output. Hence, we excluded studies that had measured traits related to fertility, such as testis size in males or egg maturation rate in females. This resulted in 12 studies for further analysis (shown in Table S6 below)

We extracted values, and the corresponding standard errors, of male and female reproductive output under benign and stressful temperature (including development at elevated temperature, heat shock and ramping temperatures). Figure 4 only includes pairs of values where the stress treatment led to a statistically significant decrease in fertility in at least one sex. Supplementary table S6 contains all treatments of the included studies, also such without significant effects in either sex. We calculated the TSF as the proportional loss in fertility due to temperature stress:

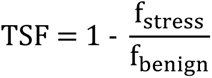

We then proceeded to calculate a standardised estimate of sexual dimorphism in TSF:

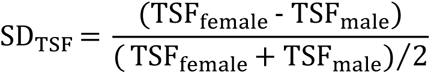

A given SD_TSF_ from this study was derived by first calculating the TSF of one sex in a given experimental condition (e.g., TSF_female_ with remating but no cohabitation from experiment *1.2*) and then always using the TSF observed in one of the two main experiments (*1.1* for juvenile stress and *2.1* for adult stress) for the opposite sex from control populations as comparison (e.g., TSF_male_ from experiment *1.1*). To assess whether the obtained values of SD_TSF_ were different from zero we propagated the uncertainty in male and female reproductive output to calculate 95% confidence intervals. In five instances one sex showed a negative load (i.e., increased fertility after heat treatment), in three of these instances the opposite sex suffered a significant loss of fertility. In these three cases the negative load was set to zero for plotting, resulting in the respective data to appear at a value of -2 or 2 (see Supplementary table S6 for all values including negative loads). We performed no formal statistical analysis on this dataset as it only serves to illustrate the comparison of intra- and interspecific variation in TSF found in this and other studies, and the number of studies is limited.

**Supplement S6:**
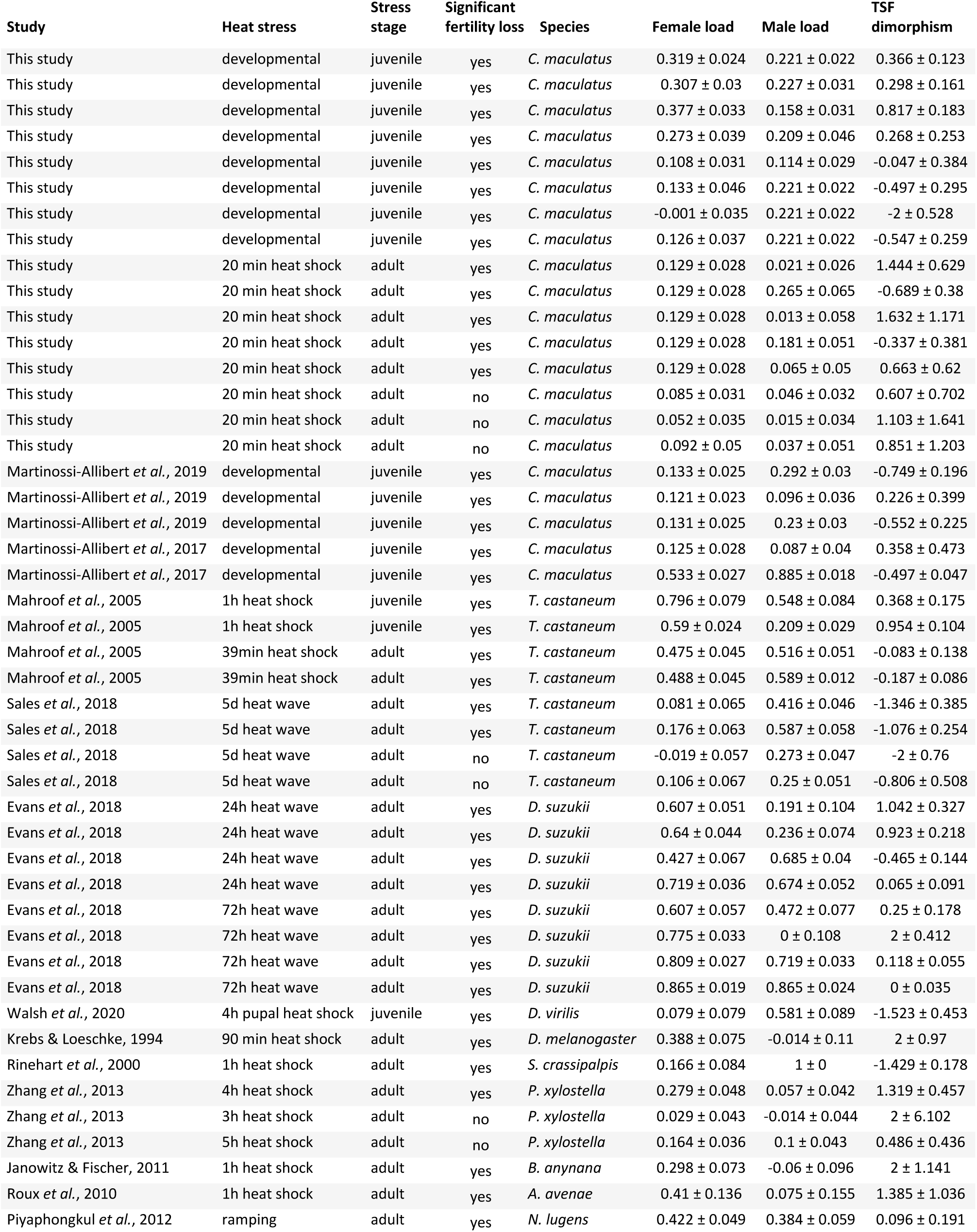
Values extracted from studies meeting the criteria outlined in supplement S5. Only values obtained for stress conditions under which at least one sex showed a significant fitness reduction were included in the figures presented in the main manuscript. (2012)

| Study | Heat stress | Stress stage | Significant fertility loss | Species | Female load | Male load | TSF dimorphism |
| --- | --- | --- | --- | --- | --- | --- | --- |
| This study | developmental | juvenile | yes | *C. maculatus* | 0.319 ± 0.024 | 0.221 ± 0.022 | 0.366 ± 0.123 |
| This study | developmental | juvenile | yes | *C. maculatus* | 0.307 ± 0.03 | 0.227 ± 0.031 | 0.298 ± 0.161 |
| This study | developmental | juvenile | yes | *C. maculatus* | 0.377 ± 0.033 | 0.158 ± 0.031 | 0.817 ± 0.183 |
| This study | developmental | juvenile | yes | *C. maculatus* | 0.273 ± 0.039 | 0.209 ± 0.046 | 0.268 ± 0.253 |
| This study | developmental | juvenile | yes | *C. maculatus* | 0.108 ± 0.031 | 0.114 ± 0.029 | -0.047 ± 0.384 |
| This study | developmental | juvenile | yes | *C. maculatus* | 0.133 ± 0.046 | 0.221 ± 0.022 | -0.497 ± 0.295 |
| This study | developmental | juvenile | yes | *C. maculatus* | -0.001 ± 0.035 | 0.221 ± 0.022 | -2 ± 0.528 |
| This study | developmental | juvenile | yes | *C. maculatus* | 0.126 ± 0.037 | 0.221 ± 0.022 | -0.547 ± 0.259 |
| This study | 20 min heat shock | adult | yes | *C. maculatus* | 0.129 ± 0.028 | 0.021 ± 0.026 | 1.444 ± 0.629 |
| This study | 20 min heat shock | adult | yes | *C. maculatus* | 0.129 ± 0.028 | 0.265 ± 0.065 | -0.689 ± 0.38 |
| This study | 20 min heat shock | adult | yes | *C. maculatus* | 0.129 ± 0.028 | 0.013 ± 0.058 | 1.632 ± 1.171 |
| This study | 20 min heat shock | adult | yes | *C. maculatus* | 0.129 ± 0.028 | 0.181 ± 0.051 | -0.337 ± 0.381 |
| This study | 20 min heat shock | adult | yes | *C. maculatus* | 0.129 ± 0.028 | 0.065 ± 0.05 | 0.663 ± 0.62 |
| This study | 20 min heat shock | adult | no | *C. maculatus* | 0.085 ± 0.031 | 0.046 ± 0.032 | 0.607 ± 0.702 |
| This study | 20 min heat shock | adult | no | *C. maculatus* | 0.052 ± 0.035 | 0.015 ± 0.034 | 1.103 ± 1.641 |
| This study | 20 min heat shock | adult | no | *C. maculatus* | 0.092 ± 0.05 | 0.037 ± 0.051 | 0.851 ± 1.203 |
| Martinossi-Allibert *et al.*, 2019 | developmental | juvenile | yes | *C. maculatus* | 0.133 ± 0.025 | 0.292 ± 0.03 | -0.749 ± 0.196 |
| Martinossi-Allibert *et al.*, 2019 | developmental | juvenile | yes | *C. maculatus* | 0.121 ± 0.023 | 0.096 ± 0.036 | 0.226 ± 0.399 |
| Martinossi-Allibert *et al.*, 2019 | developmental | juvenile | yes | *C. maculatus* | 0.131 ± 0.025 | 0.23 ± 0.03 | -0.552 ± 0.225 |
| Martinossi-Allibert *et al.*, 2017 | developmental | juvenile | yes | *C. maculatus* | 0.125 ± 0.028 | 0.087 ± 0.04 | 0.358 ± 0.473 |
| Martinossi-Allibert *et al.*, 2017 | developmental | juvenile | yes | *C. maculatus* | 0.533 ± 0.027 | 0.885 ± 0.018 | -0.497 ± 0.047 |
| Mahroof *et al.*, 2005 | 1h heat shock | juvenile | yes | *T. castaneum* | 0.796 ± 0.079 | 0.548 ± 0.084 | 0.368 ± 0.175 |
| Mahroof *et al.*, 2005 | 1h heat shock | juvenile | yes | *T. castaneum* | 0.59 ± 0.024 | 0.209 ± 0.029 | 0.954 ± 0.104 |
| Mahroof *et al.*, 2005 | 39min heat shock | adult | yes | *T. castaneum* | 0.475 ± 0.045 | 0.516 ± 0.051 | -0.083 ± 0.138 |
| Mahroof *et al.*, 2005 | 39min heat shock | adult | yes | *T. castaneum* | 0.488 ± 0.045 | 0.589 ± 0.012 | -0.187 ± 0.086 |
| Sales *et al.*, 2018 | 5d heat wave | adult | yes | *T. castaneum* | 0.081 ± 0.065 | 0.416 ± 0.046 | -1.346 ± 0.385 |
| Sales *et al.*, 2018 | 5d heat wave | adult | yes | *T. castaneum* | 0.176 ± 0.063 | 0.587 ± 0.058 | -1.076 ± 0.254 |
| Sales *et al.*, 2018 | 5d heat wave | adult | no | *T. castaneum* | -0.019 ± 0.057 | 0.273 ± 0.047 | -2 ± 0.76 |
| Sales *et al.*, 2018 | 5d heat wave | adult | no | *T. castaneum* | 0.106 ± 0.067 | 0.25 ± 0.051 | -0.806 ± 0.508 |
| Evans *et al.*, 2018 | 24h heat wave | adult | yes | *D. suzukii* | 0.607 ± 0.051 | 0.191 ± 0.104 | 1.042 ± 0.327 |
| Evans *et al.*, 2018 | 24h heat wave | adult | yes | *D. suzukii* | 0.64 ± 0.044 | 0.236 ± 0.074 | 0.923 ± 0.218 |
| Evans *et al.*, 2018 | 24h heat wave | adult | yes | *D. suzukii* | 0.427 ± 0.067 | 0.685 ± 0.04 | -0.465 ± 0.144 |
| Evans *et al.*, 2018 | 24h heat wave | adult | yes | *D. suzukii* | 0.719 ± 0.036 | 0.674 ± 0.052 | 0.065 ± 0.091 |
| Evans *et al.*, 2018 | 72h heat wave | adult | yes | *D. suzukii* | 0.607 ± 0.057 | 0.472 ± 0.077 | 0.25 ± 0.178 |
| Evans *et al.*, 2018 | 72h heat wave | adult | yes | *D. suzukii* | 0.775 ± 0.033 | 0 ± 0.108 | 2 ± 0.412 |
| Evans *et al.*, 2018 | 72h heat wave | adult | yes | *D. suzukii* | 0.809 ± 0.027 | 0.719 ± 0.033 | 0.118 ± 0.055 |
| Evans *et al.*, 2018 | 72h heat wave | adult | yes | *D. suzukii* | 0.865 ± 0.019 | 0.865 ± 0.024 | 0 ± 0.035 |
| Walsh *et al.*, 2020 | 4h pupal heat shock | juvenile | yes | *D. virilis* | 0.079 ± 0.079 | 0.581 ± 0.089 | -1.523 ± 0.453 |
| Krebs & Loeschke, 1994 | 90 min heat shock | adult | yes | *D. melanogaster* | 0.388 ± 0.075 | -0.014 ± 0.11 | 2 ± 0.97 |
| Rinehart *et al.*, 2000 | 1h heat shock | adult | yes | *S. crassipalpis* | 0.166 ± 0.084 | 1 ± 0 | -1.429 ± 0.178 |
| Zhang *et al.*, 2013 | 4h heat shock | adult | yes | *P. xylostella* | 0.279 ± 0.048 | 0.057 ± 0.042 | 1.319 ± 0.457 |
| Zhang *et al.*, 2013 | 3h heat shock | adult | no | *P. xylostella* | 0.029 ± 0.043 | -0.014 ± 0.044 | 2 ± 6.102 |
| Zhang *et al.*, 2013 | 5h heat shock | adult | no | *P. xylostella* | 0.164 ± 0.036 | 0.1 ± 0.043 | 0.486 ± 0.436 |
| Janowitz & Fischer, 2011 | 1h heat shock | adult | yes | *B. anynana* | 0.298 ± 0.073 | -0.06 ± 0.096 | 2 ± 1.141 |
| Roux *et al.*, 2010 | 1h heat shock | adult | yes | *A. avenae* | 0.41 ± 0.136 | 0.075 ± 0.155 | 1.385 ± 1.036 |
| Piyaphongkul *et al.*, 2012 | ramping | adult | yes | *N. lugens* | 0.422 ± 0.049 | 0.384 ± 0.059 | 0.096 ± 0.191 |

